# Functional hot-spots and communication networks of the cellular prion protein from information-theoretical measures

**DOI:** 10.64898/2026.11.20.768072

**Authors:** Alessia Guadagnin Pattaro, Dino Gasparotto, Emiliano Biasini, Raffaello Potestio

**Affiliations:** Physics Department, University of Trento, Trento, Italy; INFN-TIFPA, Trento Institute for Fundamental Physics and Applications, Trento, Italy; Department of Cellular, Computational and Integrative Biology, University of Trento, Trento, Italy

## Abstract

The cellular prion protein (PrP^C^) is a ∼33 kDa molecule featuring a disordered region and a globular part, mostly expressed in the brain and in the central nervous system. The biological role of this protein is still unclear, but its pathogenic behavior in specific conditions has been ascertained. In fact, for reasons that are still under scrutiny, PrP^C^may convert to its misfolded counterpart PrP^Sc^, which accumulates in tissues and gives rise to a plethora of neurodegenerative diseases. At this day, the mechanism underlying the misfolding process of PrP^C^ to PrP^Sc^ is not fully understood. Here, we analyze molecular dynamics simulations of PrP^C^ by means of information-theoretical approaches to single out specific residues that might play a key functional role in the protein, and to identify the network of communication that relays mechanical signals among distal regions of the molecule. We found such *hot-spots* in the *α*_1_helix and the *β*_2_-*α*_2_hinge, which are involved in the initiation of the misfolding process; furthermore, the pair Val209-Cys214 acts as the main link for the information relay between the two aforementioned regions. These observations suggest that the main determinants that trigger the pathogenic conformational transition of PrP^C^’s globular domain are inherently encoded in its structure; this result opens the possibility of developing therapeutic molecules that interfere with these functional elements to mitigate the protein’s misfolding and aggregation.

**Author summary:** Prion diseases are fatal neurodegenerative disorders associated with the conversion of the normal cellular prion protein into an abnormal, disease-associated form. Despite extensive research, the molecular mechanisms underlying this process remain incompletely understood, particularly how different regions of the protein interact and communicate. In this study, we used computer simulations and two complementary information-based methods to investigate the internal organization and communication pathways of the human prion protein. We identified three regions of particular importance: the *α*_1_helix, the interface between the *α*_2_and *α*_3_helices, and the flexible hinge connecting the *β*_2_strand and *α*_2_helix. We also identified a major communication pathway involving four amino acids, Tyr150, Val209, Cys214, and Tyr162, which links these regions through the highly conserved *α*_3_helix. Notably, several identified regions overlap with sites associated with disease-causing mutations. We further found that opening of the *α*_1_helix is associated with reduced fluctuations in other parts of the protein. Our findings reveal how the intrinsic structural organization of the protein supports long-range communication and suggest that alterations in these pathways may contribute to its destabilization and pathogenic conversion. These results may help identify new targets for future therapeutic strategies.

## Introduction

The cellular prion protein (PrP^C^, Fig. 1) is central to a class of fatal neurodegenerative disorders known as transmissible spongiform encephalopathies, including Creutzfeldt–Jakob disease and bovine spongiform encephalopathy [1]. These diseases are associated with the conformational conversion of PrP^C^, which is predominantly *α*-helical, into a misfolded, *β*-sheet–rich pathogenic form, PrP^Sc^ [2]. Despite extensive experimental efforts, and still limited yet significant computer-aided studies, the molecular mechanisms underlying this conversion remain incompletely understood, particularly at the level of residue-specific interactions and long-range communication within the protein [3, 4].

**Fig 1.**
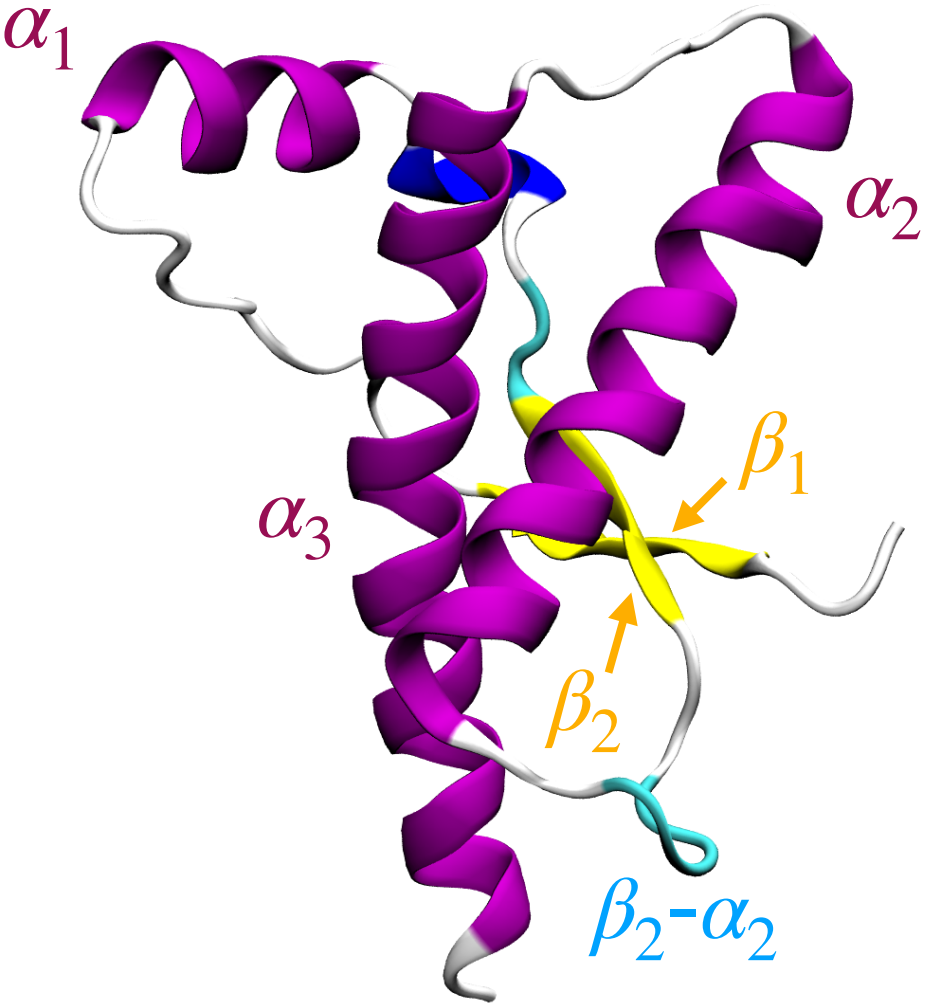
PrP^C^globular domain. Rendering of the PrP^C^globular domain colored according to its secondary structure, with its *α*-helices and *β*-sheets labeled.

A key aspect of protein function and misfunction is allosteric modulation [5–8], i.e., the propagation of local perturbations through the protein structure, ultimately affecting distant sites. In the context of PrP^C^, identifying the elements of the molecule that mediate such communication represents a promising avenue to understand both structural stability and pathological misfolding [3]. More specifically, the combination of molecular dynamics (MD) simulations with information-theoretic analysis approaches apt to identify the key structural elements that underlie the molecule’s function-oriented dynamics and support allosteric communication would return relevant insight into these questions. To date, however, only a limited number of studies have systematically investigated the intramolecular properties of PrP^C^through MD simulations at atomistic resolution [3, 4, 9–11].

In the present work, we apply a computational analysis pipeline to extensive equilibrium MD simulations of the human PrP^C^, in order to identify the most relevant residues and reconstruct the principal communication routes within the molecule. To this end, we employ two approaches: the mapping entropy optimization workflow, or MEOW for short, and the mutual information communication network. The MEOW method [12, 13] enables the identification of important, putatively functional residues of a protein in a totally unsupervised manner, without relying on prior biological assumptions about the system. The second method consists in the construction of a communication graph between residue pairs, with links weighted through the normalized mutual information (NMI) [14]; this graph can inform about long (sequence, spatial) distance in the molecule and highlight highly correlated albeit distal residues.

Our results highlight specific regions and amino acids that act as central hubs in the protein, and suggest mechanistic links between these and known structural features associated with prion conversion and disease.

## Materials and methods

In this work we analyzed the five MD simulations of the globular part of PrP^C^obtained by Spagnolli *et al*. [15], for a total simulation time of 21 *µ*s. As anticipated, our investigation relies on two techniques: the mapping entropy optimization workflow (MEOW) [12, 13], and the normalized mutual information (NMI) [14, 16]. The first method, developed by some of us, allows us to identify subgroups of *N* residues that optimally convey information about the conformational and energetic variability of the whole protein. By computing the empirical probability of a residue to be part of such optimal subgroups, the method assigns to it a measure of its structural, energetic, and ultimately functional importance, or *relevance*. Previous applications of the MEOW approach, in fact, have shown that the identified hot-spots are majorly involved in the function of the protein under examination [12, 17], thereby supporting and corroborating the correlation between this information-theoretic importance and biological significance.

The second measure is the normalized mutual information; this is employed in the method, implemented by Doering *et al*. [14] in the package MDPath, that evaluates the amount of information that is shared between two residues. This pair-wise quantity is then employed in reconstructing the putative communication network that spans throughout the whole protein, thereby allowing us to infer which areas of the structure are important for signal propagation between distal regions. The obtained graph is then analyzed in order to retrieve the hubs through which the majority of information is relayed.

The five equilibrium MD simulations of the human prion protein, kindly provided by M. Rigoli, were produced and investigated in a study by Spagnolli and coworkers [15]. The mapping entropy optimization was carried out independently on the five trajectories; the relevance for each atom was calculated as the probability for an atom to appear in a mapping, while the corresponding relevance value for an amino acid was taken as the relevance of its most conserved atom. Since the optimization process is stochastic, 48 independent runs were performed for each trajectory to obtain a reliable statistics of residue relevance. The calculation was performed varying the number *N* of atoms retained in the optimal mappings; in particular, we considered a number of sites equal to the number of C_*α*_ atoms (*N* = 102), then 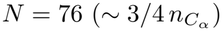, 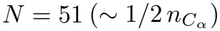, and 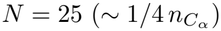. The results of this analysis are reported in Fig. 2, where one can appreciate those amino acids that are consistently reported as being relevant (e.g. Arg156, Gln172).

**Fig 2.**
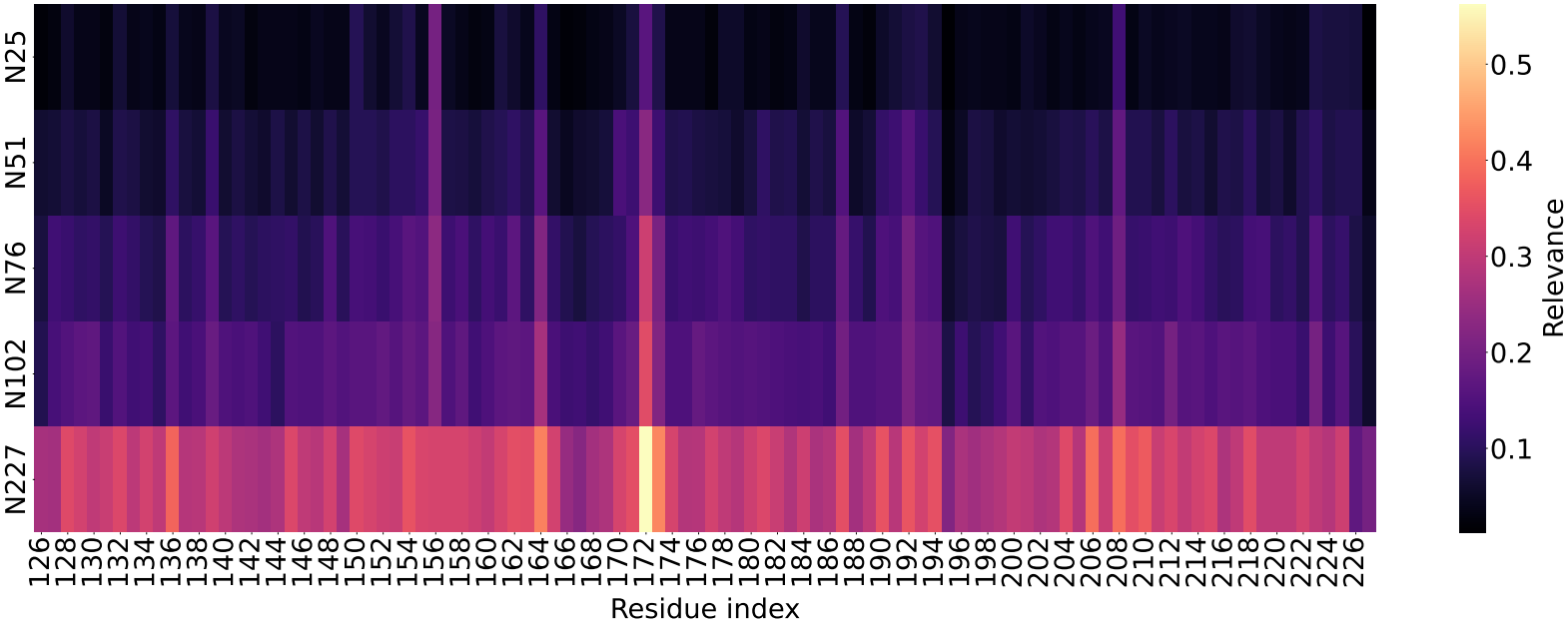
Varying retained sites heatmap. Heatmap of the aggregated MEOW relevance data for different numbers *N* of retained sites. The number *N* increases from the bottom row to the top row.

To assess the robustness of the optimization procedure, and hence of the relevance values, we compared the results obtained applying MEOW to the five trajectories independently *versus* a single aggregated trajectory, in all cases using *N* = 102. The outcome of this analysis, reported in Fig S1 and Fig. S2, supports the statistical significance of the identified relevant residues. Additionally, the MEOW results obtained from the MD trajectories were compared against a null model of the protein that retains realistic local fluctuation dynamics but lacks any intramolecular correlation: this comparison corroborates the noteworthiness of the relevant residues. Details and data about this analysis are reported in Fig. S9.

As for the NMI network, the information pathways were identified independently on the five trajectories using the software MDPath by Doering *et al*. [14]. The method calculates normalized mutual information NMI of the dihedral angles to reconstruct the putative communication pathways. In this case, the NMI-weighted graph generated by the software was used to retrieve the most significant routes by considering the 20 edges with the highest weight using the Python library Networkx [18], which was also used to perform graph centrality analyses. The aggregated results were obtained by considering the top 20 edges weighted by NMI of each graph to create an *intersection graph*, with the new weights *w* being the number of times each edge appeared in the five trajectories’ list of top 20 nodes. Only edges present in at least two trajectories have been considered. The results for the single trajectories are reported in SI (Figs.S3 to S7).

Finally, we performed a study of the root mean square fluctuations (RMSF) of the C_*α*_ atoms of the molecule in two distinct conformational basins, identified on the basis of a quantitative descriptor of the open/closed state of the *α*_1_loop. More detailed about this analysis are provided hereafter.

## Results and Discussion

The globular part of the cellular prion protein (PrP^C^) can be divided into two subdomains: the first – comprising the *β*_1_, *α*_1_, and *β*_2_ chains – features higher fluctuations, while the second, constituted by the remaining *α*_2_ and *α*_3_ chains, is intrinsically stable and the least flexible region of the protein [4, 19, 20]. These two subdomains are joined by the flexible hinge *β*_2_-*α*_2_ [15, 21–25]. The residues singled out as the most relevant by the MEOW approach in the *N* = 25 case (see Materials and Methods, Fig. 2) are centered around specific areas of PrP^C^: the aforementioned *β*_2_-*α*_2_ hinge with the amino acid Gln172, Asn173, and Arg164, the *α*_1_helix with Tyr150, Met154, and Arg156; the top part of *α*_2_ and *α*_3_ with Thr190 and Thr193; and finally the core of the *α*_3_ helix with Arg208 and Gln223. These results are illustrated in Fig. 3.

**Fig 3.**
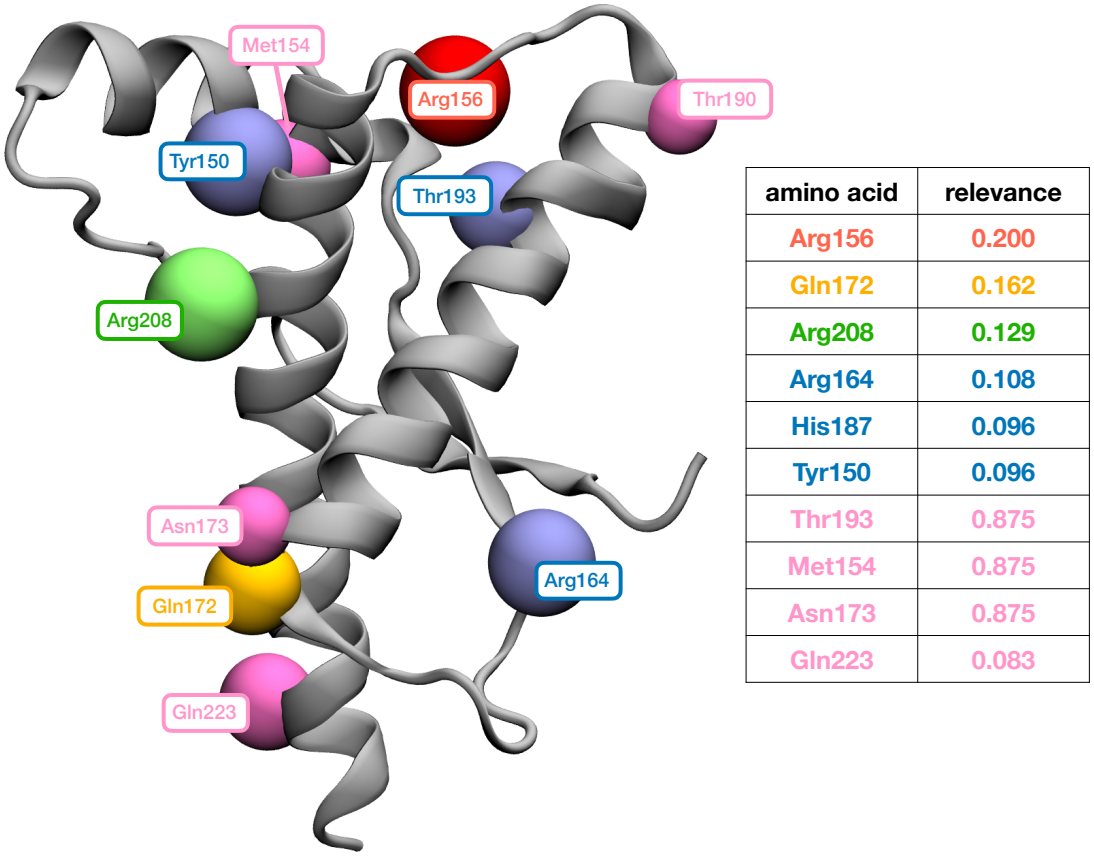
MEOW-E results for PrP^C^. Graphical representation of PrP^C^with the location and the label of the ten most relevant amino acids according to MEOW (aggregated results of the five independent trajectories) with *N* = 25 retained sites (see Materials and Methods). The different label colors indicate the ranking of the amino acids according to the relevance.

The communication network graph obtained through the NMI analysis, instead, is reported in Fig. 4, with the most important edges colored according to the aggregated NMI strength, and the amino acid nodes they connect represented as colored spheres. The resulting graph has four components, the largest of which is shown by itself in Fig. 4(a). Within the graph, the edge with the largest weight is the amino acid pair Val209–Cys214; this also happens to be the edge with the largest betweenness centrality [26], that is, the probability that an edge or a node appears in the shortest path between two other nodes. This pair thus seems to act as the major information relay within PrP^C^. Notably, both residues involved in this edge are located in the structurally autonomous and highly conserved *α*_3_ helix [19], which, together with *α*_2_, exhibits relatively low backbone flexibility [20]. In particular, Cys214 has been identified as a highly conserved residue displaying low conformational flexibility and contributing to stabilize the structural core through the disulfide bridge connecting *α*_2_ and *α*_3_. Furthermore, Val209 belongs to an *α*_3_ segment involved in prominent correlated dynamic motions with the C-terminal portion of the same helix [20]. These observations suggest that the identified edge involves structurally constrained residues potentially contributing to the dynamical communication network of the protein.

**Fig 4.**
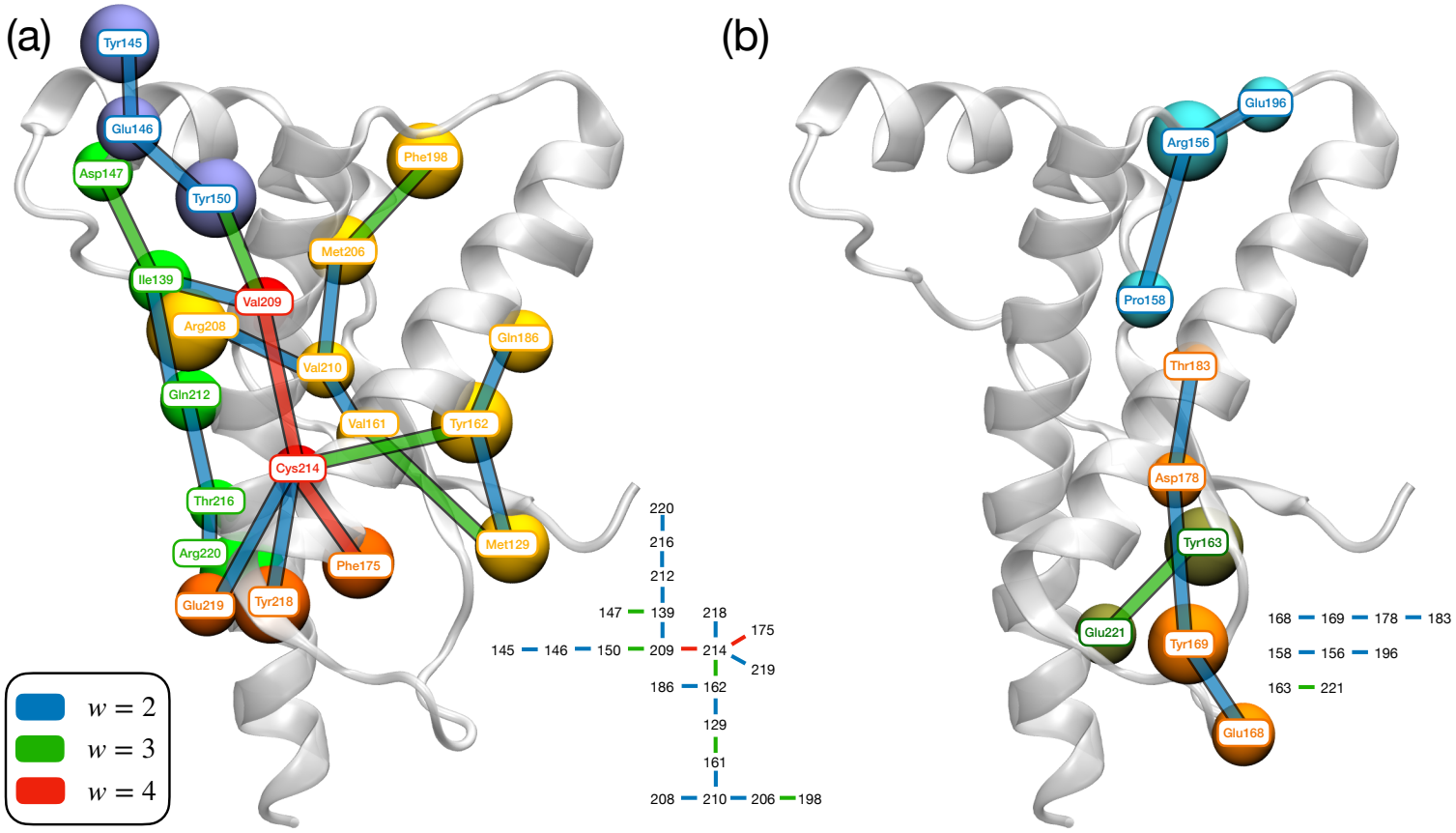
MI network results for PrP^C^. Graphical and schematic representation of the NMI graph obtained with MDPath [14]. The edges of the protein graphs and the schematic graphs (at the bottom) are colored according to the number of independent trajectories they appeared in (legend at bottom left). (a) Main network, the amino acids in red are the central information core of the protein. (b) Secondary networks and isolated pair.

The other edges with high betweenness help us understand the information flow. From Val209 we reach Tyr150, which is located on *α*_1_, while from Cys214 we go to Tyr162 and to Phe175, respectively at the beginning and at the end of the *β*_2_-*α*_2_ loop. The information flow axis suggests that the dynamics of the *α*_1_ helix and the *β*_2_-*α*_2_ hinge, despite being distant in the structure, are tightly intertwined. Most importantly, the communication passes through the highly conserved *α*_3_ helix. From the axis Tyr150–Val209–Cys214–Tyr162, the information branches across the whole PrP^C^structure. Some distal connections exist also between the *β*_2_-*α*_2_ hinge and the final part of *α*_2_(see Fig. 4(b)).

The results of the MEOW relevance and of the NMI graph share important, coherent characteristics: in fact, the majority of the hot-spots are located along the main communication axis. In greater detail, we can partition PrP^C^in three main areas of interest: (a) *α*_1_ and the *β*_1_-*α*_1_ loop, which attains an open conformation only in a single instance of the MD simulations [15]. (b) The area enclosed by *α*_1_ and the top part of *α*_2_ and *α*_3_ [27–30], which has been shown to be a drug binding site during the folding process of PrP^C^; we will refer to this region as the intermediate folding (IF) binding site. Note that for this area to act as a binding site the protein has to be in a conformation where the *α*_1_ helix is open [15]. (c) The already mentioned *β*_2_-*α*_2_ hinge, which is important for promoting the conversion of PrP^C^to its misfolded, pathogenic counterpart PrP^Sc^, especially regarding the presence of mutations in residues 170–175 [22], but also during the misfolding mechanism, as shown by Sabzehei *et al*. [4]. According to this recent work, the globular part of PrP^C^needs to unfold first, then to adapt to the PrP^Sc^template; the unfolding process is initiated by the separation of the more fluctuating *α*_1_-*β*_1_-*β*_2_ subregion from the rigid *α*_2_-*α*_3_ using the *β*_2_-*α*_2_ loop as a *door hinge*. In light of these findings, the high relevance of *β*_2_-*α*_2_, of *α*_3_, and of the *α*_1_ helices, along with the strong communication axis that connects the three areas, advocates for a model in which perturbations, as in disease-associated mutations occurring in this region, might affect local stability or disrupt the global information relay, with potentially impactful consequences for the misfolding process to PrP^Sc^. The same considerations are valid also for the IF binding pocket, which is connected to the *β*_2_-*α*_2_ hinge in the network (Fig. 4b) and entail some residues that are identified as hot-spots of PrP^C^.

The role of the *α*_2_–*α*_3_ interface is even more important when considering the PrP glycosylation. PrP^C^is N-glycosylated at two asparagine residues, Asn181 and Asn197, both located in close spatial proximity to the *α*_2_ helix and, consequently, near the pocket region [31]. Although glycosylation is not explicitly modeled in the present study, its presence is expected to modulate the local environment of the *α*_2_–*α*_3_ interface through steric effects, altered hydration patterns, and changes in inter-helical interactions. In particular, glycans attached at Asn181 and Asn197 may influence the accessibility and conformational flexibility of the proposed drug-binding pocket, potentially stabilizing specific conformational substates or shielding key residues from solvent exposure [32]. From a network perspective, such modifications could perturb the communication pathways identified in this work by altering residue–residue correlations in this region. This is especially relevant given that glycosylation patterns are known to affect prion strain properties and disease progression. The overlap between glycosylation sites and the hot-spot residues in the *α*_2_–*α*_3_ interface therefore suggests a possible mechanistic link in which post-translational modifications modulate the communication within the protein and, in turn, influence the propensity of PrP^C^ to undergo pathogenic conversion.

As anticipated, in the course of the MD simulations a single event of *α*_1_ opening was registered, a movement that allows PrP^C^ to be in a conformation similar to the folding intermediate hosting the binding pocket [15]. According to Sabzehei *et al*. , in the unfolding process leading to the formation of PrP^Sc^ the first part that opens up is *α*_1_, followed by *β*_1_ and *β*_2_ [4]; hence, we explored whether the opening movement is associated to conformations in which the other regions of the molecule fluctuate more, possibly hinting at a greater structural instability and ultimately to the unfolding of *β*_1_ and *β*_2_. To this end, we grouped the sampled conformations in an open and a closed state, employing the distance between the C_*α*_’s of His140 and Arg208 to discriminate between them. Since the open conformation appeared only briefly (about 0.484 *µ*s, 425 frames), the sampling of the two basins is largely inhomogeneous (Fig. S8). Keeping this in mind, we calculated the RMSF of the C_*α*_ atoms of the two states, reported in Fig. 5: the open conformation features very intense fluctuations at the N-terminal and a small bump at the hinge region, while the rest of the protein does not show any significant signal; the closed conformation, in contrast, has a higher RMSF at both terminals, as well as at the *β*_2_-*α*_2_ hinge and in the part of *α*_2_ belonging to the IF binding site. The most fluctuating areas that we found are consistent with those involved in the first steps of the unfolding process identified by Sabzehei *et al*. [4], but the fact that there are greater fluctuations in the closed state is thought-provoking: the opening of the *α*_1_ helix seems to favor a stiffening of the structure instead of its relaxation, while in the closed state the hinges fluctuate more broadly. This is also confirmed by the average values of RMSF for the full structure and the two hinges, as reported in Table 1.

**Table 1.** Average values of RMSF for the open and close conformations of the whole PrP^C^structure, of the *α*_2_-*α*_3_, and of the *β*_2_-*α*_2_hinges.

|  |  | **avg RMSF (Å)** |
|---|---|---|
| PrP<sup>C</sup> | open | 1.8 |
|  | close | 2.5 |
| α<sub>2</sub>-α<sub>3</sub> | open | 1.3 |
|  | close | 4.3 |
| β<sub>2</sub>-α<sub>2</sub> | open | 1.8 |
|  | close | 3.5 |

**Fig 5.**
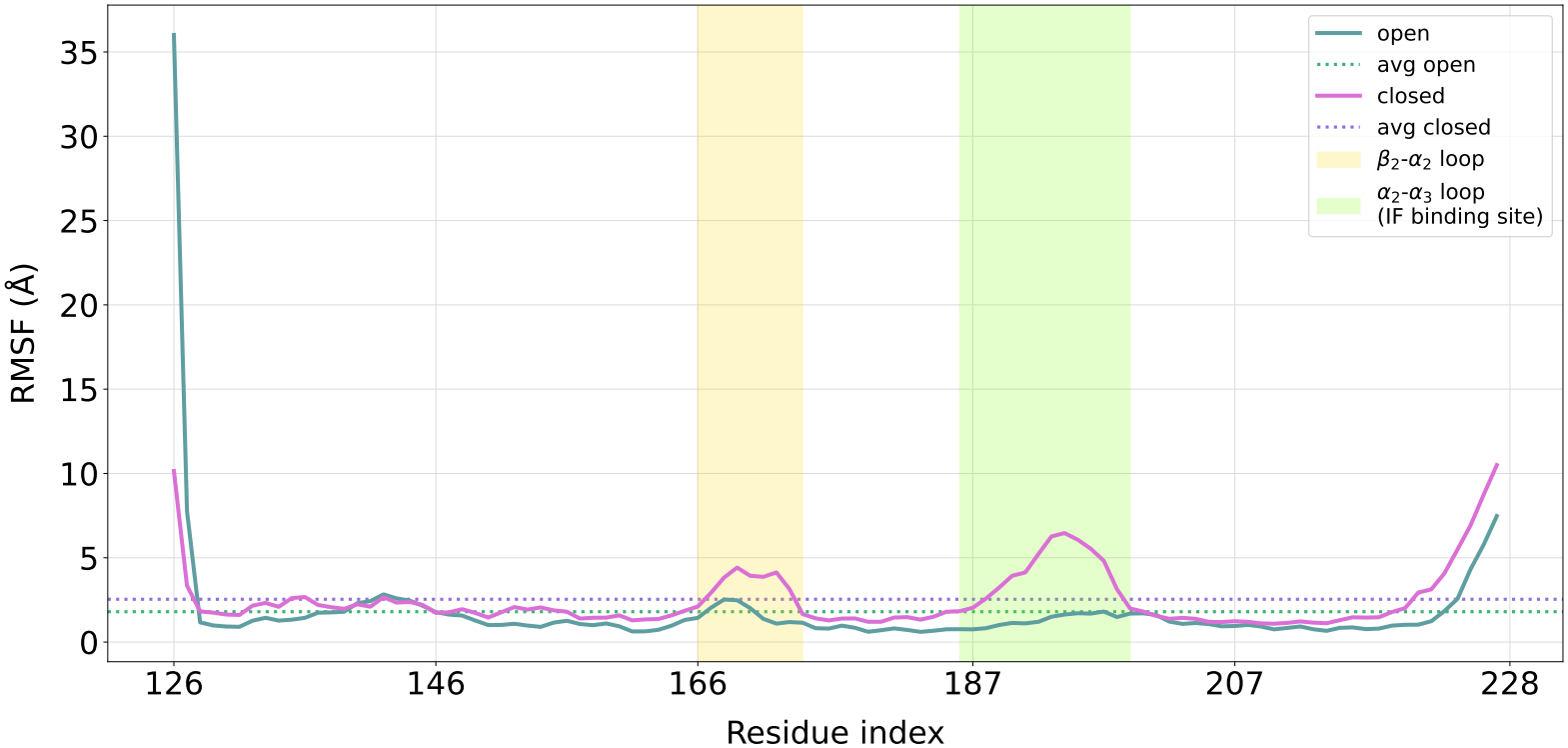
RMSF of the C_*α*_ atoms of PrP^C^. RMSF of the C_*α*_ atoms of PrP^C^in the open (green) and close (pink) conformations. The *β*_2_-*α*_2_ loop is highlighted in yellow, while the *α*_2_-*α*_3_ is highlighted in green.

## Conclusion

While the pathogenic role of PrP^C^, and in particular of its misfolded counterpart PrP^Sc^, has been clearly and rigorously established, the details of the mechanisms through which this protein gives rise to a number of diseases are still largely obscure. In order to shed light on PrP^C^function, *in silico* simulations and the analysis of their trajectories can be of valuable help.

Here, two different yet complementary methods, the mapping entropy optimization workflow (MEOW) and the normalized mutual information (NMI) network analysis, have been applied to the investigation of extensive MD simulations of the PrP^C^ globular domain. The results of this study offer a coherent picture of the internal organization of PrP^C^: in particular, the hot-spot residues with the highest MEOW relevance, namely Gln172, Arg164, and Arg208, and the main communication axis Tyr150-Val209-Cys214-Tyr162, indicate that PrP^C^ dynamics mainly involves the *α*_1_ and *α*_3_ helices and the *β*_2_-*α*_2_ hinge. Several pathogenic mutations are located at the hot-spots or near MEOW-relevant regions: for instance, D178N [33], which is associated with familial Creutzfeldt–Jakob disease (fCJD) and fatal familial insomnia (FFI), occurs in close proximity to the *β*_2_–*α*_2_ loop, the same area where highly ranked residues like Gln172 and Asn173 are located. Similarly, mutations in the *α*_2_–*α*_3_ region, including V210I [28] and E200K (one of the most common causes of fCJD [33]), directly involve residues or structural environments highlighted by our analysis. The prominence of Val210 as a key node in the communication network is particularly striking in this context, as its mutation is known to significantly alter disease susceptibility [28]. The disruption of these communication routes, whether through mutations, environmental changes, or ligand binding, may represent a key step in the loss of native stability, and favor the conformational transition to misfolded forms. The correspondence between the identified hubs and known disease-associated mutations further suggests that these regions are critical determinants of prion pathogenicity, providing a mechanistic basis for the insurgence of disease, and offering potential targets for therapeutic intervention. Importantly, the agreement between the present findings and previous studies is achieved despite the fully unsupervised nature of our information-theoretic approach, which does not incorporate prior knowledge of PrP^C^biology: the identified residues and pathways, which reflect intrinsic dynamical properties of PrP^C^, overlap with known pathogenic mutation sites identified through independent, experimental means. This correlation provides an *a posteriori* validation of the method and underscores its potential for identifying functionally and clinically relevant regions to exploit in the development of pharmaceutical strategies to mitigate the impact of prion-related diseases.

## Supporting information

**S1 MEOW-E aggregated trajectories** Heatmap reporting the aggregated results for MEOW-E for the five indepentent trajectories for *N* = 102.

**S2 MEOW-E separated trajectories** Heatmaps reporting the results for MEOW-E for the five trajectories one by one.

**S3 MI pathways for t_1** Graphical and schematic depiction of the results for the MI pathways of t_1. On the schematic graphs (bottom left and right), the top 10 edges according to NMI are depicted in magenta, edges from 11 to 20 in blue. (a) Main network. (b) Secondary connections.

**S4 MI pathways for t_2** Graphical and schematic depiction of the results for the MI pathways of t_2. On the schematic graphs (bottom left and right), the top 10 edges according to NMI are depicted in magenta, edges from 11 to 20 in blue. (a) Main network. (b) Branching network from Cys214. (c) Lone pairs.

**S5 MI pathways for t_3** Graphical and schematic depiction of the results for the MI pathways of t_3. On the schematic graphs (bottom left and right), the top 10 edges according to NMI are depicted in magenta, edges from 11 to 20 in blue. (a) Main network. (b) Branching network from Cys214.

**S6 MI pathways for t_4** Graphical and schematic depiction of the results for the MI pathways of t_4. On the schematic graphs (bottom left and right), the top 10 edges according to NMI are depicted in magenta, edges from 11 to 20 in blue. (a) Main network. (b) Branching network from Val210. (c) Secondary connections and lone pair.

**S7 MI pathways for t_5** Graphical and schematic depiction of the results for the MI pathways of t_5. On the schematic graphs (bottom left and right), the top 10 edges according to NMI are depicted in magenta, edges from 11 to 20 in blue. (a) Main network and branching networks for Asp147, Val210, and Gln212. (b) Secondary connection and lone pair.

**S8 Distance distribution between residues 140-208** Histogram reporting the frame distribution after the separation of the five PrP MD trajectories according to the distance between the C_*α*_ atoms His140 and Arg208. The separation threshold, highlighted in yellow, is chosen at_15 Å.

**S9 Comparison between PrP**^**C**^**trajectories and PrP independent model** Panel reporting the results of the mapping optimization and of the random extraction of mappings for the five PrP trajectories (top, MEOW-E results in green) and for the independent model (bottom, MEOW-E results in blue), each one with the respective 95% confidence interval. The independent model is an elastic network model [34, 35] whose terms of the Hessian matrix have been nullified, so the topology of the protein is retained but without any cross-interaction between the residues. The independent model was built with the ProDy 2.0 python library [36] while the random mapping extraction has been carried on using EXCOGITO [13].

## Acknowledgments

The authors wish to thank Marta Rigoli and Jesús R. Requena for a critical reading and insightful comments.

This work was supported by grants from Fondazione Telethon (GGP20043, GMR24T2072), the American CJD Foundation and the Italian Association for Prion Encephalopathies (AIENP) to E.B.

This work was conducted in the spirit of the Slow Science Manifesto, advocating for collaborative and sustainable research (slow-science.com).

## Data and software availability

Raw data produced and analyzed in this work are freely available on the Zenodo repository doi.org/10.5281/zenodo.24336655.

