## Supplementary material for "Functional hot-spots and communication networks of the cellular prion protein from information-theoretical measures"



(Dated: November 20, 2026)



*

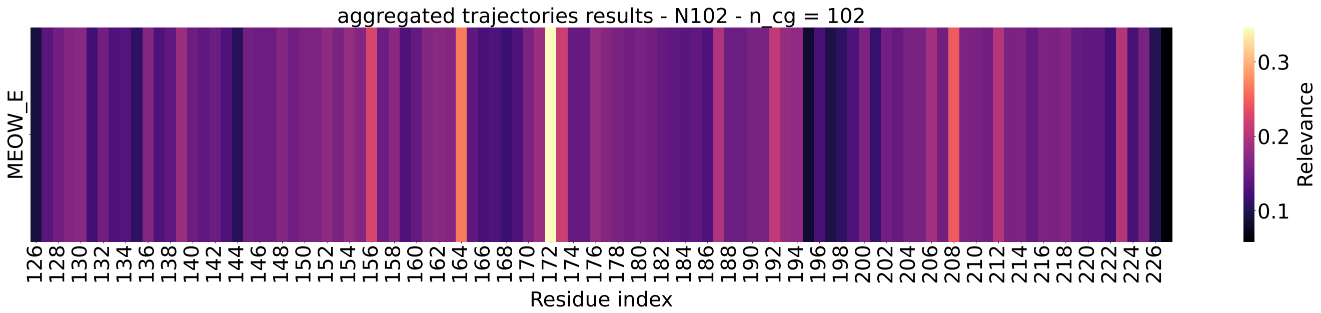


Figure 1: Heatmap reporting the aggregated results for MEOW-E for the five indepentent trajectories.

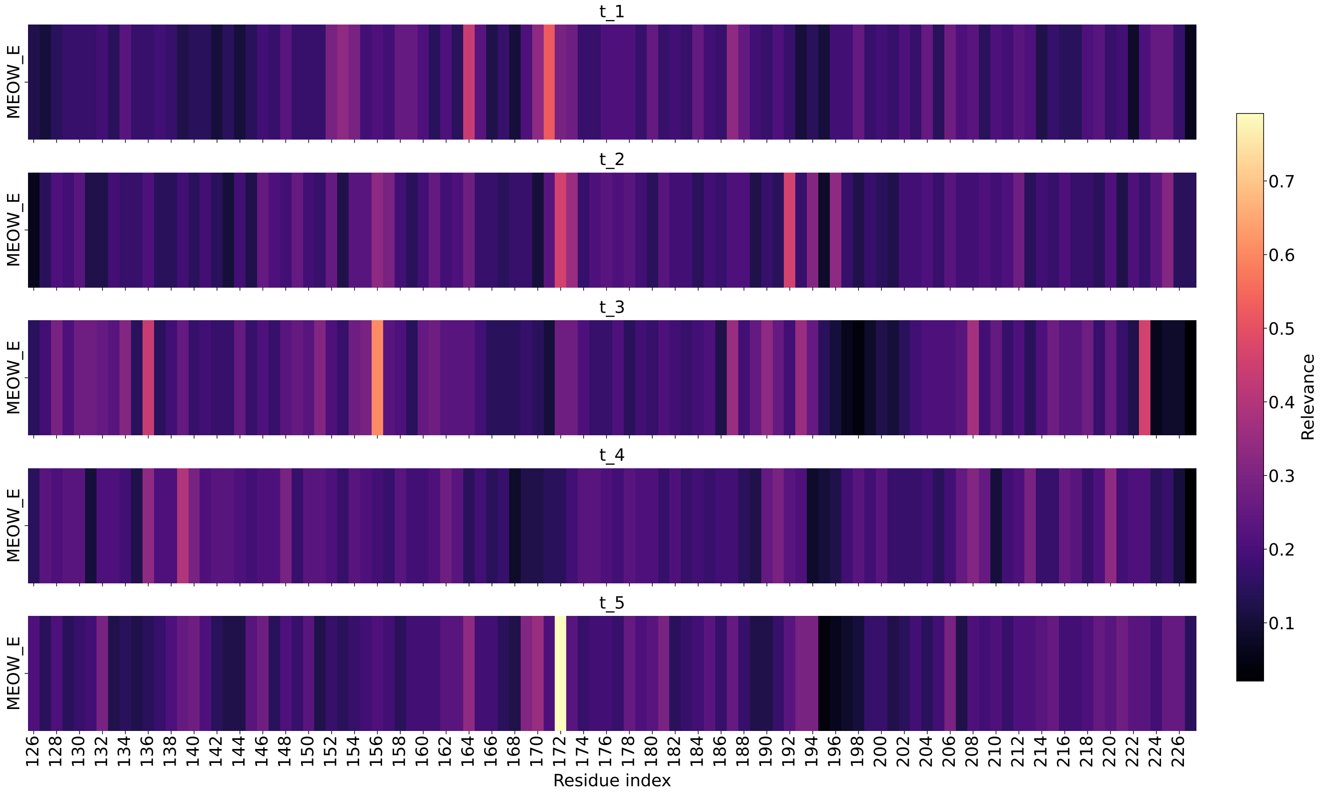


Figure 2: Heatmaps reporting the results for MEOW-E for the five trajectories one by one.

[1] M. M. Tirion, "Large amplitude elastic motions in proteins from a single-parameter, atomic analysis," *Physical review letters*, vol. 77, no. 9, p. 1905, 1996.

[2] X.-Q. Yao, L. Skjærven, and B. J. Grant, "Rapid characterization of allosteric networks with ensemble normal mode analysis," *The journal of physical chemistry. B*, vol. 120, no. 33, p. 8276, 2016.

[3] S. Zhang, J. M. Krieger, Y. Zhang, C. Kaya, B. Kaynak, K. Mikulska-Ruminska, P. Doruker, H. Li, and I. Bahar, "Prody 2.0: increased scale and scope after 10 years of protein dynamics modelling with python," *Bioinformatics*, vol. 37, no. 20, pp. 3657–3659, 2021.

[4] M. Giulini, R. Fiorentini, L. Tubiana, R. Potestio, and R. Menichetti, "Excogito, an extensible coarse-graining toolbox for the investigation of biomolecules by means of low-resolution representations," *Journal of Chemical Information and Modeling*, vol. 64, no. 12, pp. 4912–4927, 2024.

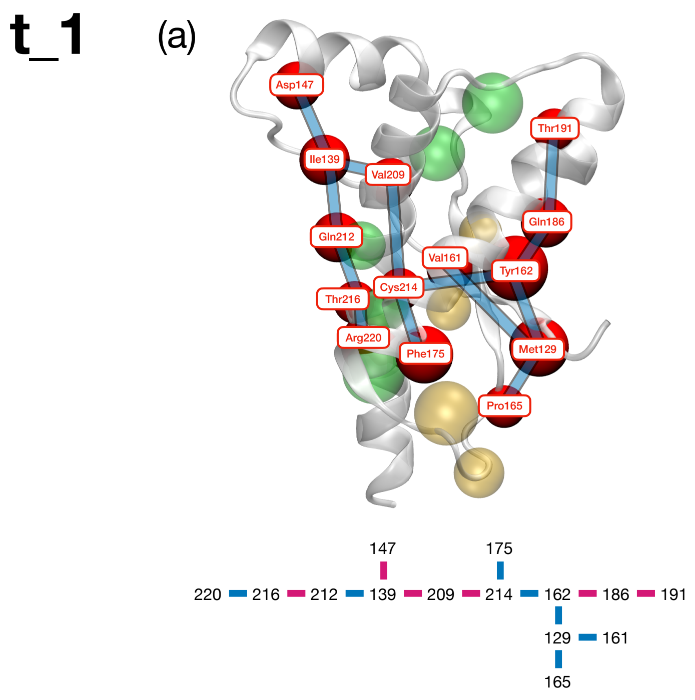


Figure 3: Graphical and schematic depiction of the results for the MI pathways of t_1. On the schematic graphs (bottom left and right), the top 10 edges according to NMI are depicted in magenta, edges from 11 to 20 in blue. (a) Main network. (b) Secondary connections.

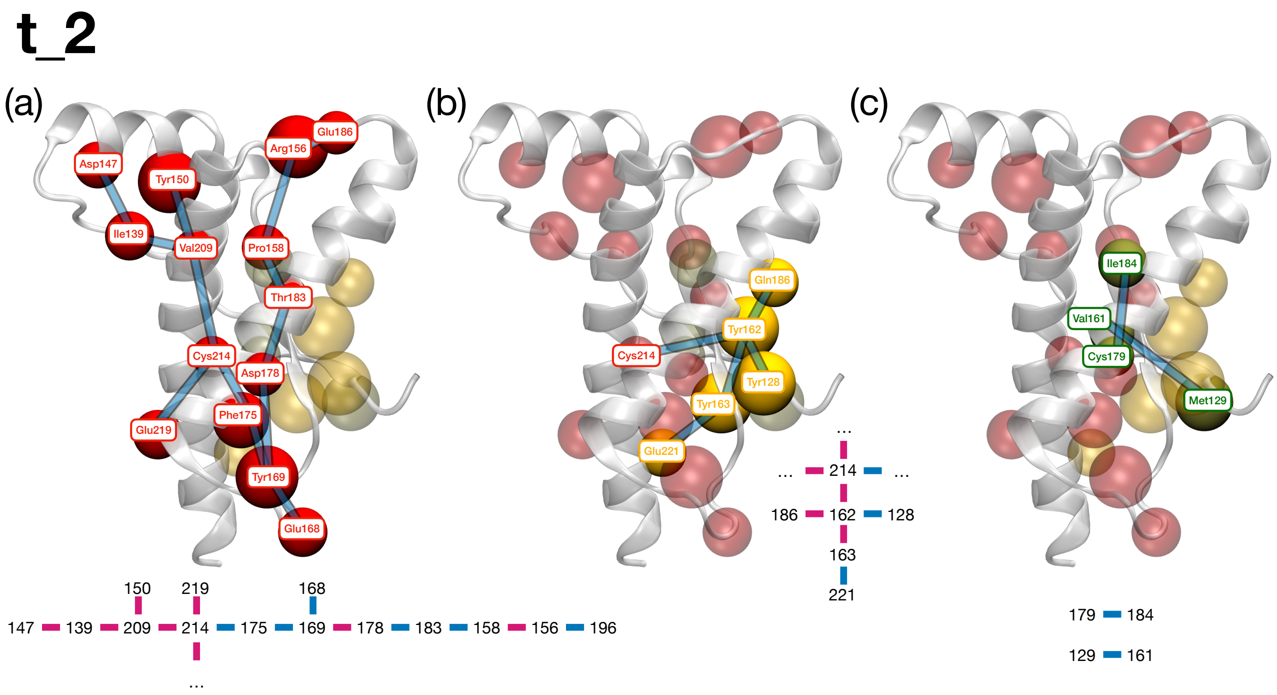


Figure 4: Graphical and schematic depiction of the results for the MI pathways of t_2. On the schematic graphs (bottom left and right), the top 10 edges according to NMI are depicted in magenta, edges from 11 to 20 in blue. (a) Main network. (b) Branching network from Cys214. (c) Lone pairs.

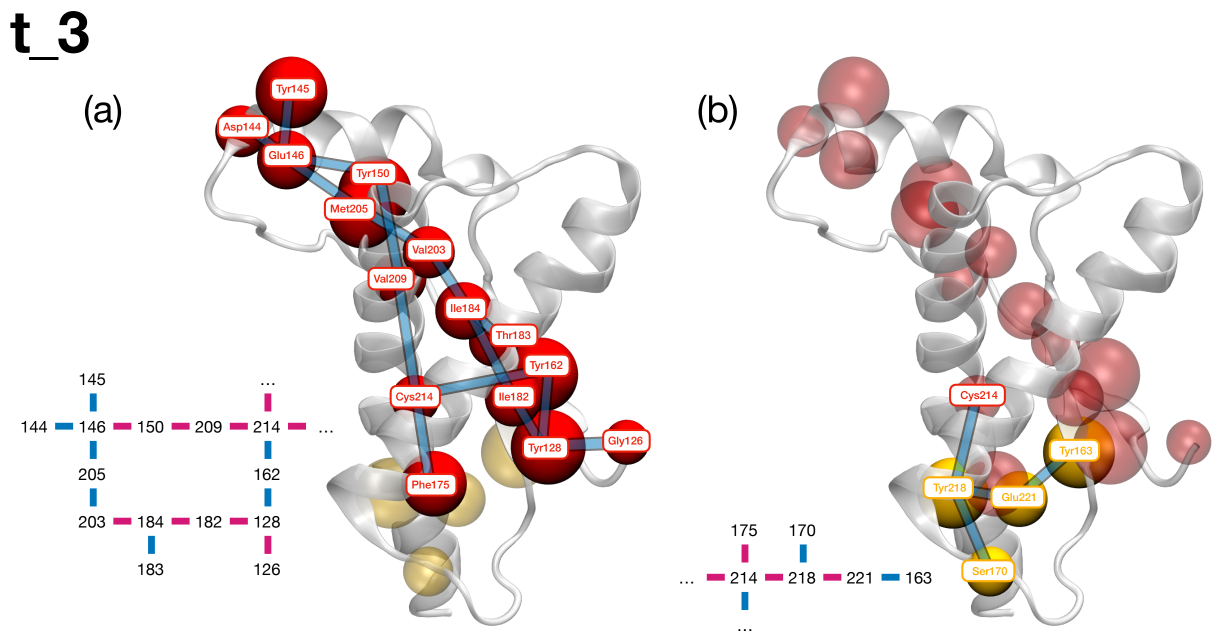


Figure 5: Graphical and schematic depiction of the results for the MI pathways of t_3. On the schematic graphs (bottom left and right), the top 10 edges according to NMI are depicted in magenta, edges from 11 to 20 in blue. (a) Main network. (b) Branching network from Cys214.

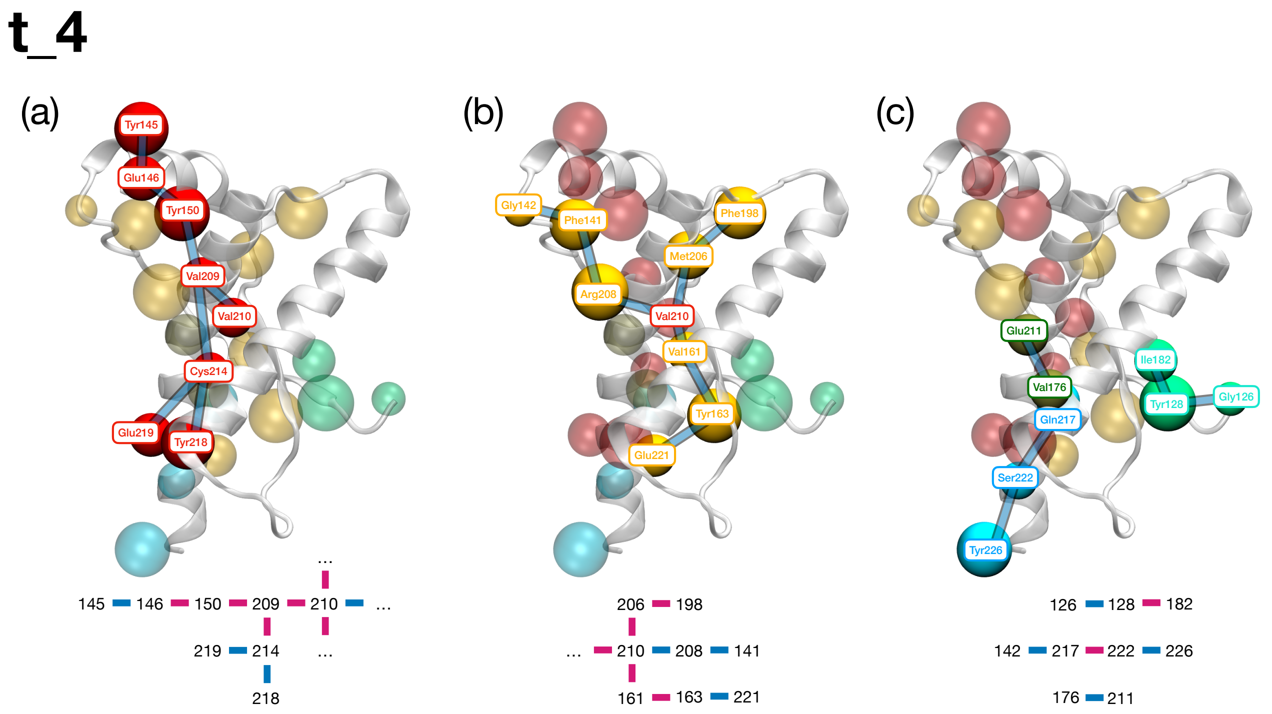


Figure 6: Graphical and schematic depiction of the results for the MI pathways of t_4. On the schematic graphs (bottom left and right), the top 10 edges according to NMI are depicted in magenta, edges from 11 to 20 in blue. (a) Main network. (b) Branching network from Val210. (c) Secondary connections and lone pair.

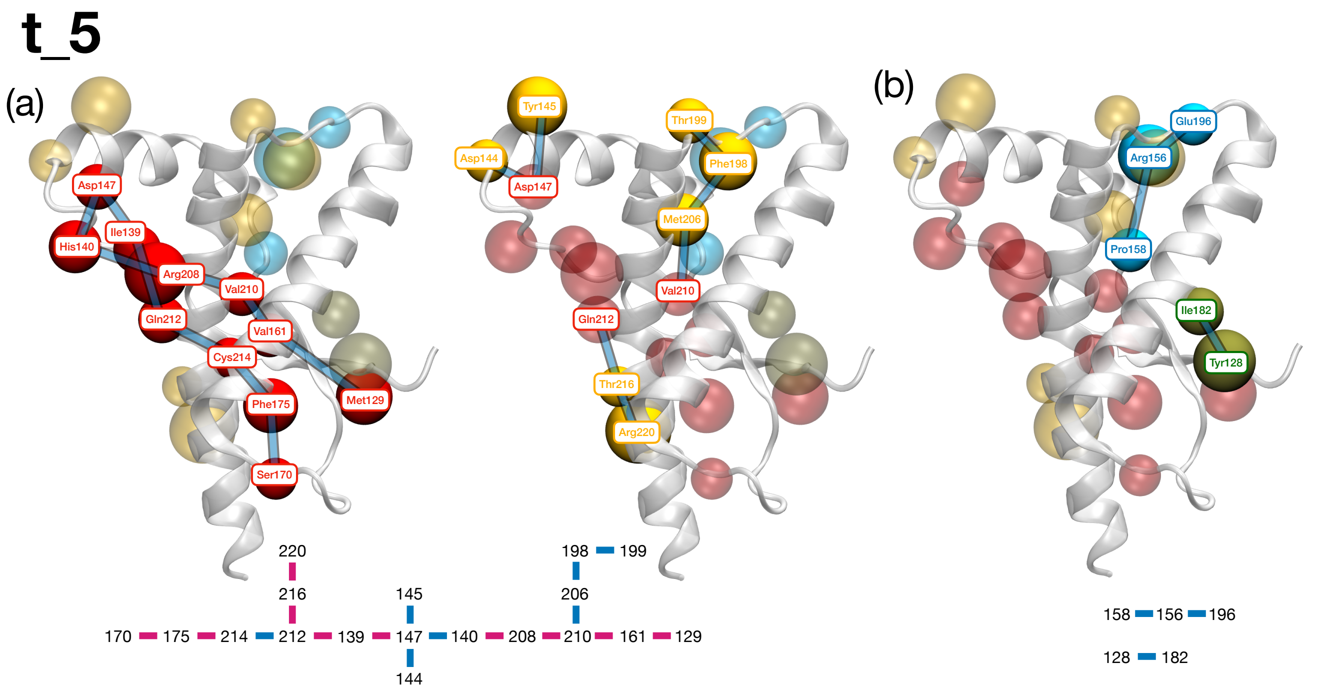


Figure 7: Graphical and schematic depiction of the results for the MI pathways of t_5. On the schematic graphs (bottom left and right), the top 10 edges according to NMI are depicted in magenta, edges from 11 to 20 in blue. (a) Main network and branching networks for Asp147, Val210, and Gln212. (b) Secondary connection and lone pair.

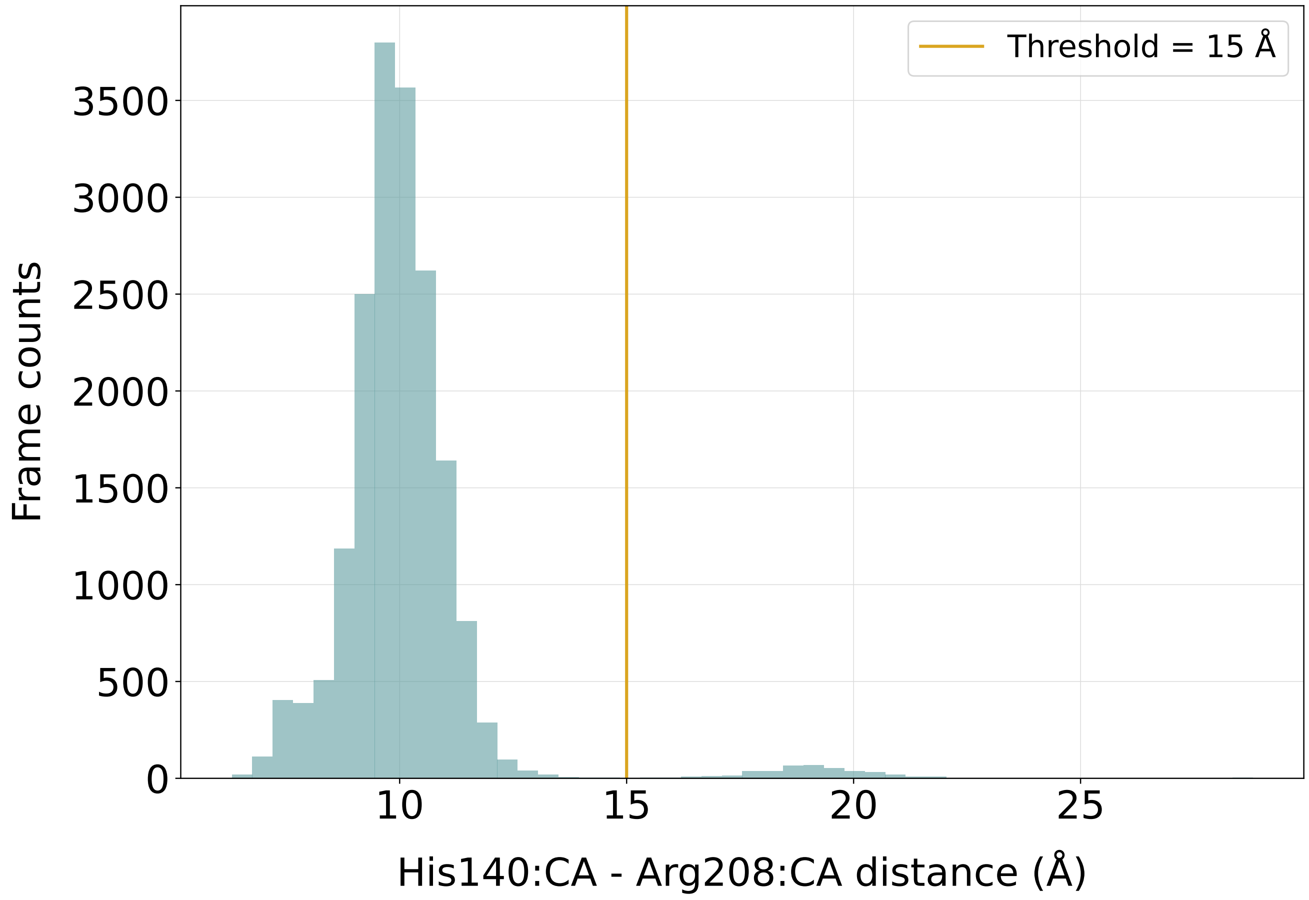


Figure 8: Histogram reporting the frame distribution after the separation of the five PrP MD trajectories according to the distance between the $C_\alpha$ atoms His140 and Arg208. The separation threshold, highlighted in yellow, is chosen at 15 Å.

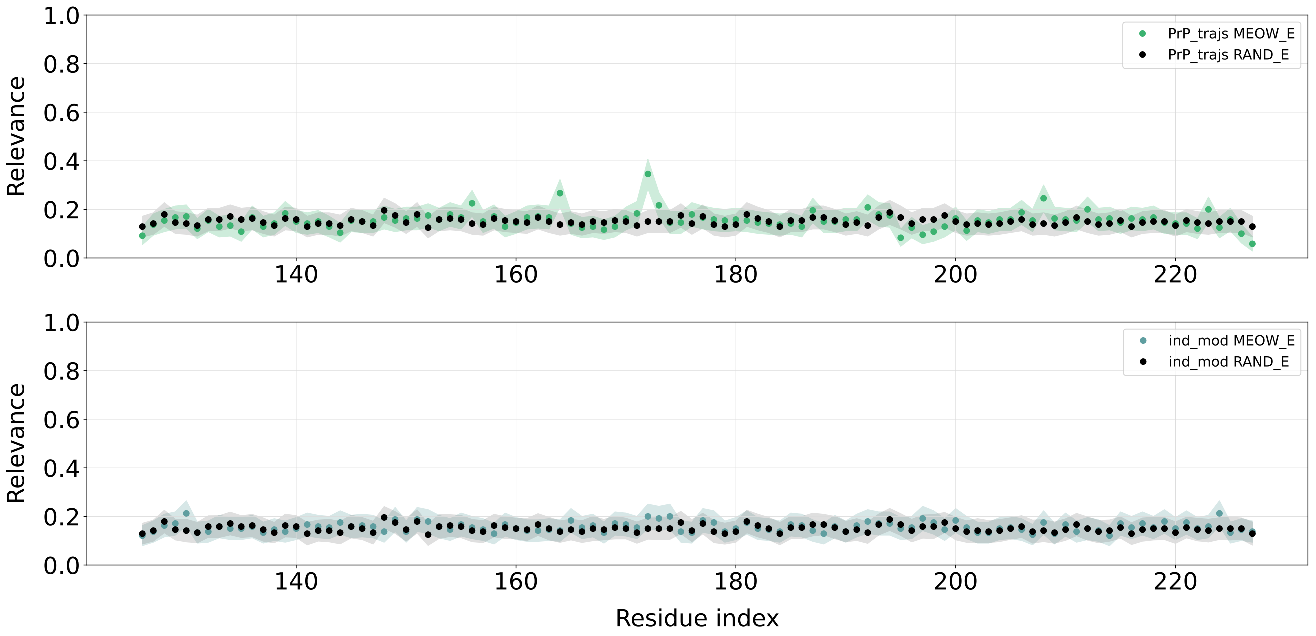


Figure 9: Panel rep orting the results of the mapping optimization and of the random extraction of mappings for the five PrP trajectories (top, MEOW-E results in green) and for the independent model (bottom, MEOW-E results in blue), each one with the respective 95% confidence interval. The independent model is an elastic network model [1, 2] whose terms of the Hessian matrix h ave been nullified, so the topology of the protein is retained but without any cross-interaction between the residues. The independent model was built with the ProDy 2.0 python library [3] while the random mapping extraction has been carried on using EXCOGITO [4].
